# Using the ANDROMEDA by Prosilico Software for Prediction of the Human Pharmacokinetics of 4 Compounds of Natural Origin - Colistin, Curucumin, UCN-01 and Voclosporin

**DOI:** 10.1101/2022.08.17.504228

**Authors:** Urban Fagerholm, Sven Hellberg, Jonathan Alvarsson, Ola Spjuth

## Abstract

**Background:** It is important that pharmacokinetic (PK) prediction methods are validated, and also for compounds with varying physicochemical properties, molecular weights and PK characteristics.

**Methods:** The objective was to investigate how well the ANDROMEDA by Prosilico software predicts the clinical PK of four compounds of natural origin and with PK obstacles, not yet fully characterized PK, and/or inaccurate lab method-based predictions - colistin (negligible absorption, good metabolic stability, significant excretion), curucumin (low solubility, apparently poor bioavailability), UCN-01 (extremely high degree of plasma protein binding, metabolic stability, long half-life and poor PK prediction) and voclosporin (poorly understood PK).

**Results:** All categorial predictions except one were correct, and the median prediction error was 2.5-fold. Largest prediction errors were found for the unbound fraction in plasma (>24-fold), clearance (178-fold) and half-life (90-fold) of UCN-01. Corresponding errors for clearance and half-life obtained with allometry were greater, 5800- and 145-fold, respectively. Extremely high affinity for alpha1-acid glycoprotein could explain these large prediction errors for this compound. A substantial amount of data and knowledge was added with the predictions.

**Conclusion:** Despite challenging compounds and PK, predictions were comparably good. The results further validated ANDROMEDA by Prosilico for human clinical PK-predictions.

## Introduction

We have developed and validated software for prediction for human clinical pharmacokinetics (PK) - ANDROMEDA by Prosilico (1–4). The software system is based on conformal prediction (CP), which is a methodology that sits on top of machine learning methods and produce valid levels of confidence (5), unique algorithms and a new human physiologically-based pharmacokinetic (PBPK) model (1). For a more extensive introduction to CP we refer to Alvarsson et al. 2014 (6).

In a validation study with pharmaceutical drugs, the Q^2^ (forward-looking predictive accuracy) for log volume of distribution (V_ss_) was 0.65, and the mean prediction error was 2.4-fold (2). 69 % of test compounds had an observed V_ss_ within the prediction interval at a 70 % confidence level, which empirically validated the model (it predicts at the selected confidence level) (2).

It is important that prediction methods are validated and investigated for different types of compounds, for example, for compounds with varying physicochemical properties, molecular weight (MW) and PK-characteristics.

In a first attempt to challenge ANDROMEDA by Prosilico, the human PK of chemicals of various classes was predicted and compared to observed data (3). In this validation study, median and mean prediction errors for the PK-parameters V_ss_, fraction absorbed (f_a_), oral bioavailability (F), half-life (t_½_), unbound fraction in plasma (fu), clearance (CL) and fraction excreted (fe) were 1.3- to 2.7-fold and 1.4- to 4.8-fold, respectively (3).

ANDROMEDA by Prosilico was then further tested/challenged through predictions of the f_a_ of macrocyclic drugs, which have MW above its major domain (MW 100-700 g/mole) (4). A 2-fold median prediction error for f_a_ was reached (4).

Among drugs there are compounds of natural origin with complex structures and PK and relatively high MW. Examples include colistin (natural compound), curucumin (natural compound), UCN-01 (synthetic derivative of natural staurosporine) and voclosporin (analogue of the natural drug cyclosporin). PK challenges/obstacles, not yet fully characterized PK, and/or inaccurate lab method-based PK-predictions make them interesting to use in validation studies of human clinical PK-prediction methods.

The cyclic polypeptidic antibiotic colistin (MW 1155 g/mole; log P=2.4) is characterized by high aqueous solubility (564 g/L), negligible gastrointestinal absorption (no clinically useful f_a_ and F), low CL (26 mL/min), good metabolic stability (approximated intrinsic metabolic CL (CL_int_) of 27 mL/min), significant excretion (63 % of unchanged colistin excreted renally following an intravenous dose), intermediate f_u_ (0.34), small V_ss_ (0.09 L/kg) and long t½ (18.5 h according to a recent study) in man, and adhesion to lab materials (including plastics) (7–9). It is unknown whether colistin is excreted unchanged via bile.

Curucumin (MW=368 g/mole; log P=3.6) is a highly lipophilic (predicted log P=3.6) phenolic compound (the major yellow pigment extracted from the spice turmeric) with poor aqueous solubility (5.8 mg/L). Its PK is characterized by absorption into the systemic circulation mainly or solely as conjugates (gram doses required to detect curcumin in blood; f_a_ in rats = 60 %), crossing of the blood-brain barrier, approximated poor oral F (0.5-1 % F in rats) and extensive metabolism (its phenolic structure indicates that it is likely to undergo significant extraction in the gut wall mucosa) (10–14). The fate of the compound during absorption from the gastrointestinal tract and into the blood stream and its distribution and elimination are not fully understood.

UCN-01 (MW=483 g/mole; log P=2.4) is a permeable and highly soluble (predicted 79 mg/L) anticancer compound (protein kinase inhibitor) with an extremely high specific affinity for alpha1-acid glycoprotein (AAG) (f_u_<0.02 %), low V_ss_ (0.11 L/kg, extremely low CL (0.18 mL/min), CL_int_ > 1000 mL/min and extremely long t_½_ (ca 1 month) (15–19). In contrast, its V_ss_ and CL are high in mice, rats and dogs (15). Allometric scaling (animal-to-man predictions) of V_ss_ and CL resulted in 63- and 5800-fold overpredictions, respectively (15,19). Based on this established methodology, a 145-fold underprediction of t½ and ca 1 million-fold underprediction of the systemic exposure (potential health risk) is expected following oral dosing to humans.

The cyclic polypeptide voclosporin (MW 1215 g/mole; log P=8.6), marked in 2021, is a calcineurin inhibitor used as an immunosuppressant medication. It has a f_u_ of 3 % and t_½_ of 30 h (20). Much of its PK-characteristics in man is unknown.

The objective of the study was to evaluate how well ANDROMEDA by Prosilico predicts the human clinical PK of the pharmacokinetically challenging compounds colistin, curucumin, UCN-01 and voclosporin.

## Methods

### Compounds and Clinical Data

Colistin (MW=1155 g/mole; log P=-2.4; aqueous solubility=564 g/L), curucumin (MW=368 g/mole; log P=3.6; aqueous solubility=5.8 mg/L), UCN-01 (MW=483 g/mole; log P=2.4; aqueous solubility=79 mg/L) and voclosporin (MW 1215 g/mol; log P=8.6; aqueous solubility <100 mg/L) were selected as test compounds. For these compounds there was a limited amount of available clinical PK data and information - only 10, 2, 4 and 5 parameter estimates for colistin, curucumin, UCN-01 and voclosporin, respectively.

### ANDROMEDA by Prosilico

The prediction system based on CP and the new PBPK-model (ANDROMEDA software by Prosilico for prediction, simulation and optimization of human clinical PK) predicts the following human PK-parameters – passive permeability class (P_e_-class), f_a_, f_a_ from colon (f_a,col_), dissolution potential (f_diss_), extended release (ER)-potential, absorption rate constant (k_a_), f_u_, f_u_ in blood (f_u,bl_), blood-to-plasma concentration ratio (C_bl_/C_pl_), passive blood-brain barrier (BBB) P_e_-class, BBB-efflux, fraction bound to brain tissue (f_b,brain_), V_ss_, CL_int_, hepatic CL (CL_H_), biliary CL (CL_bile_), renal CL (CL_R_), CL, fraction escaping gut-wall extraction (F_gut_), f_e_, oral F, t_½_, and MDR-1-, BCRP-, MRP2-, CYP2C9-, CYP2D6- and CYP3A4-specificities (1–4).

ANDROMEDA by Prosilico is mainly applicable for compounds with molecular weight 100 to 700 g/mole. There are groups of compounds for which its *in silico* models do not work, including metals and quaternary amines, and have limited use, for example, hydrolysis sensitive compounds and drugs binding covalently and/or to DNA. For a more extensive introduction to ANDROMEDA by Prosilico see Fagerholm et al. (1–4).

None of the selected compounds in the present study was included in the training sets of the used CP models. Thus, every prediction was a forward-looking prediction where each compound in the benchmarking set was unknown to the models.

Predicted and measured/observed data were compared both numerically and categorically (low/moderate/high). Upper:lower limits for moderate f_a_, F, f_u_, f_e_, CL, CL_H_, CL_R_, CL_int_, V_ss_ and t_½_ were set to 0.20:0.80, 0.20:0.80, 0.05:0.40, 0.20:0.50, 250:750 mL/min, 250:750 mL/min, 30:100 mL/min, 500:10000 mL/min, 0.5:3.0 L/kg and 3:8 h, respectively.

## Results

It was possible to predict the complete human clinical PK for the four compounds. Predictions and observations/measurements are shown in Table 1. All except one of categorial predictions (low/moderate/high; yes/no) were correct (20 out of 21). The median prediction error was 2.3-fold (n=18). Largest prediction errors were found for the f_u_ (>24-fold), CL (178-fold) and t_½_ (90-fold) of UCN-01.

**Table 1.** Observed and predicted human clinical PK of cucucumin and colistin.

|  | MW [g/mole]<br>Unit | Curucumin |  | Colistin |  |
| --- | --- | --- | --- | --- | --- |
|  |  | 368 |  | 1155 |  |
|  |  | Predicted | Observed | Predicted | Observed |
| Oral bioavailability (F) | fraction | 0.15* | Poor* | 0.007* | Ca 0* |
| Half-life ($t_{1/2}$) | [h] | 8.0* | 6.5* | 40* | 18.5* |
| Clearance (CL)** | [mL/min] | 75 | - | 15* | 26* |
| Hepatic clearance (CL<sub>H</sub>) | [mL/min] | 70 | -- | 1* | 9* |
| Intrinsic hepatic clearance (CL<sub>int</sub>) | [mL/min] | 4295 | -- | 9* | 27* |
| Renal clearance (CL<sub>R</sub>) | [mL/min] | 1.0 | - | 14* | 17* |
| Renal fraction excreted (f<sub>eR</sub>) | fraction | 0.01 | - | 0.93* | 0.63* |
| Biliary clearance (CL<sub>bile</sub>) | [mL/min] | 3.5 | - | 0.03 | - |
| Volume of distribution at steady-state (V<sub>ss</sub>) | [L/kg] | 0.48 | - | 0.48* | 0.09* |
| Fraction absorbed (f<sub>a</sub>) | fraction | 0.79 | - | 0.007* | Low* |
| Unbound fraction in plasma (f<sub>u</sub>) | fraction | 0.017 | - | 0.11* | 0.34* |
| Passive permeability-class |  | High | - | Low* | Low* |
| Dissolution potential (f<sub>diss</sub>) | fraction | 0.96 | - | 0.56 | - |
| Fraction absorbed from colon (f<sub>a,col</sub>) | fraction | 0.42 | - | 0.03 | - |
| Extended release potential |  | Mod | - | Low | - |
| Absorption rate constant (k<sub>a</sub>) | [1/h] | 1.0 | - | 0.1 | - |
| Volume of distribution in elim. phase (V<sub>dβ</sub>) | [L/kg] | 0.74 | - | 0.74 | - |
| Unbound fraction in blood (f<sub>u,bl</sub>) | fraction | 0.016 | - | 0.095 | - |
| Blood-to-plasma concentration ratio (C<sub>bl</sub>/C<sub>pl</sub>) | fraction | 0.61 | - | 0.67 | - |
| Passive blood-brain barrier permeabilit-class |  | High* | CNS active* | Low | - |
| Fraction bound to brain tissue (f<sub>b,brain</sub>)-class | fraction | Mod | - | High | - |
| Fraction excaping gut-wall extraction (F<sub>gut</sub>) | fraction | 0.20 | - | 1.00 | - |
| Fraction excreted (f<sub>e</sub>) | fraction | 0.06 | - | 0.93 | - |
| MDR-1-substrate |  | Yes | - | Yes | - |
| BCRP-substrate |  | No | - | No | - |
| MRP2-substrate |  | No | - | Yes | - |
| Blood-brain barrier-efflux |  | Yes | - | Yes | - |
| CYP2C9-substrate |  | Yes | - | Ud | - |
| CYP2D6-substrate |  | No | - | Ud | - |
| CYP3A4-substrate |  | Yes | - | No | - |

\*Prediction and observation/measurement in same category.
Ud = undetermined.

**Table 2.** Observed and predicted human clinical PK of UCN-01 and voclosporin.

|  | MW [g/mole] | UCN-01 |  | Voclosporin |  |
| --- | --- | --- | --- | --- | --- |
|  |  | 483 |  | 1215 |  |
|  | Unit | Predicted | Observed | Predicted | Observed |
| Oral bioavailability (F) | fraction | 0.57 | Mod/High | 0.08 | - |
| Half-life ($t_{1/2}$) | [h] | 8.7* | 779* | 12.1* | 30* |
| Clearance (CL)** | [mL/min] | 32* | 0.18* | 598* | 1060* |
| Hepatic clearance (CL<sub>H</sub>) | [mL/min] | 31* | Max 0.18 | 33 | - |
| Intrinsic hepatic clearance (CL<sub>int</sub>) | [mL/min] | 6745 | >1000 | 330 | - |
| Renal clearance (CL<sub>R</sub>) | [mL/min] | 0.8 | - | 9 | - |
| Renal fraction excreted (f<sub>eR</sub>) | fraction | 0.03 | - | 0.21 | - |
| Biliary clearance (CL<sub>bile</sub>) | [mL/min] | 0 | - | 2 | - |
| Volume of distribution at steady-state (V<sub>ss</sub>) | [L/kg] | 0.22* | 0.11* | 0.52 | - |
| Fraction absorbed (f<sub>a</sub>) | fraction | 0.7754 | - | 0.08 | - |
| Unbound fraction in plasma (f<sub>u</sub>) | fraction | 0.0047* | <0.0002* | 0.10 | 0.03 |
| Passive permeability-class |  | High | - | Mod | - |
| Dissolution potential (f<sub>diss</sub>) | fraction | 0.95 | - | 0.51 | - |
| Fraction absorbed from colon (f<sub>a,col</sub>) | fraction | 0.40 | - | 0.04 | - |
| Extended release potential |  | Mod | - | Low | - |
| Absorption rate constant (k<sub>a</sub>) | [1/h] | 1.0 | - | 0.2 | - |
| Volume of distribution in elim. phase (V<sub>dβ</sub>) | [L/kg] | 0.34 | - | 0.80 | - |
| Unbound fraction in blood (f<sub>u,bl</sub>) | fraction | 0.0046 | - | 0.09 | - |
| Blood-to-plasma concentration ratio (C<sub>bl</sub>/C<sub>pl</sub>) | fraction | 0.58 | - | 0.67 | - |
| Passive blood-brain barrier permeability-class |  | High | - | Mod | - |
| Fraction bound to brain tissue (f<sub>b,brain</sub>)-class | fraction | High | - | High | - |
| Fraction escaping gut-wall extraction (F<sub>gut</sub>) | fraction | 0.77 | - | 0.96 | - |
| Fraction excreted (f<sub>e</sub>) | fraction | 0.03 | - | 0.26 | - |
| MDR-1-substrate |  | Ud | - | Yes | - |
| BCRP-substrate |  | Yes | - | No* | No* |
| MRP2-substrate |  | Yes | - | Yes | - |
| Blood-brain barrier-efflux |  | Yes | - | Yes | - |
| CYP2C9-substrate |  | Yes | - | No | - |
| CYP2D6-substrate |  | No | - | No | - |
| CYP3A4-substrate |  | Yes | - | Yes* | Yes* |

\*Prediction and observation/measurement in same category; CL/F reported instead of CL for voclosporin.
Ud = undetermined.

According to the predictions, colistin has low passive intestinal and BBB P_e_, limited f_diss_ and virtually zero f_a_ and F (in line with results in animals and man). Voclosporin was predicted to have moderate passive intestinal and BBB P_e_, limited f_diss_ and low f_a_ and F, while curucumin and UCN-01 were predicted to have high passive permeability (P_e_) and f_diss_ and moderate f_a_ (consistent with 60 % uptake of curucumin in rats). A predicted oral F of 15 % for curucumin is also consistent with approximated poor oral F in and humans and rats. Colistin, curucumin, UCN-01 and voclosporin were predicted to be substrates for at least one of the major intestinal efflux and BBB-efflux transporters.

The predicted high passive BBB P_e_ and moderate fb,brain of curucumin is consistent with CNS uptake and distribution *in vivo.*

The compounds were predicted to have low V_ss_ (0.2-0.5 L/kg). Observed estimates for colistin and UCN-01 were both 0.1 L/kg. Curucumin, UCN-01 and voclosporin were predicted to have low f_u_, whereas colistin was predicted to have moderate f_u_. There was, however, a >24-fold overprediction of f_u_ for UCN-01.

Colistin was predicted to have very low CL_int_ (9 mL/min), CL_H_ (1 mL/min) and CL (15 mL/min). Observed low CL_int_ (27 mL/min), CL_H_ (9 mL/min) and CL (26 mL/min) were in line with predicted values for this compound. It could not be verified whether colistin is a substrate for CYPs 2C9, 2D6 and 3A4. Curucumin was predicted to be a substrate for CYP2C9 and CYP3A4 and have low F_gut_ (0.2; metabolism by CYP3A4 and conjugation at phenol groups). CYP2C9- and CYP3A4-substrate specificity and some gut-wall extraction was also predicted for UCN-01. In line with clinical measurements, the CL_H_ and CL of UCN-01 were low. However, there was a 178-fold underprediction of CL. Voclosporin was predicted to have a CL/F of 598 mL/min, which is close to the measured estimate in man (1060 mL/min), and to be a substrate for CYP3A4 (consistent with *in vitro*-measurement).

The four compounds were predicted to be eliminated renally (93 % for the hydrophilic colistin, 1 % for curucumin, 3 % for UCN-01 and 21 % for voclosporin). The predicted feR for colistin (0.93) was similar to the measured clinical estimate (63 %). Colistin, curucumin and voclosporin were predicted to be eliminated via bile to some minor extent and to undergo enterohepatic circulation.

The t_½_ of colistin, curucumin and voclosporin were well predicted (1.2- to 2.5-fold errors), but that of UCN-01 was underpredicted by a f_a_ctor of 90.

## Discussion

Apparently, more than 80 % of the human PK data was missing. These missing data was now be added with the *in silico* predictions, thus, adding valuable knowledge.

Examples of new information that predictions produced include bile excretion of colistin, curucumin and voclosporin and that the low oral F of curucumin appears to mainly be due to low F_gut_ (0.2) rather than limitations in *in vivo* P_e_ or hepatic extraction. At low doses curucumin is predicted to have near complete *in vivo* dissolution potential (fdiss). It has an aqueous solubility that is higher than for acyclovir, fenofibrate, lenalidomide and ziprasidone, which have more or less complete *in vivo* dissolution in man. At higher doses, such as those given clinically, poor solubility/dissolution is anticipated to contribute to bioavailability limitations for curucumin (a f_diss_ of 0.96 at low dose indicates that most of a gram oral dose can remain undissolved).

All except one of the 21 cases where predicted and observed estimates could be compared showed correct classifications. Limits for low/moderate/high is of course subjective. We tested a selection of different limits and found 19 out of 21 correct classifications and 2 predictions in the adjacent class.

The low median prediction error (2.3-fold) is consistent with our other studies where the methodology was validated (2,3). The maximum prediction errors found in the present study are significantly greater than 15-fold maximum errors in our previous validation studies. An explanation to the large prediction errors for f_u_ (>24 fold), CL (178-fold) and t_½_ (90-fold) of UCN-01 is its extremely high specific affinity for AAG and low f_u_ (the prediction methodology appears less accurate at extremely low and high values; (2)). The prediction errors for CL and t_½_ were mainly due to the underlying overprediction of f_u_. Corresponding prediction errors for CL and t_½_ obtained with allometric scaling were, however, greater, 5800- and 145-fold, respectively (15,18). The allometric method predicted high CL and V_ss_ and moderate t_½_ of UCN-01 (15,19), which is opposite to clinical observations. Our *in silico* methodology correctly predicted low CL and V_ss_ and long t_½_ for this compound. As UCN-01 is permeable, highly soluble and has very low CLH according to experiments, moderate to high f_a_ and F is assumed. This is in line with predicted f_a_ and F of 78 and 57 %, respectively. According to allometric predictions (CLH≈liver blood flow), near zero oral F is expected in humans.

It is worth noting that for compounds with very low a 200-fold lab variability has been found for f_u_ (21), which shows that there is uncertainty for any predictive system when plasma protein binding is very high. This should be considered when predicting human clinical PK and setting doses for first human trials for such compounds.

Colistin was the compound with the most available clinical data. Despite that it has a MW that is outside of the main prediction domain of our prediction system (1155 *vs* 700 g/mole) and the complex structure (cyclic polypeptide) its human PK was generally well predicted with ANDROMEDA by Prosilico. Overall good predictions were also found for the other cyclic polypeptide, voclosporin, which has a similar MW.

## Conclusion

ANDROMEDA by Prosilico predicted the human clinical PK of the four pharmacokinetically challenging compounds comparably good, with the exception for UCN-01 which is extremely strongly bound to plasma proteins. Despite large prediction errors for UCN-01 *in silico* predictions were more accurate than with allometry. *In silico* predictions also added a substantial amount of data and knowledge. Apparently, this is the first successful attempt to predict the human clinical PK for colistin, curucumin and voclosporin. The results further validate ANDROMEDA by Prosilico.

